# DIA-NN: Neural networks and interference correction enable deep coverage in high-throughput proteomics

**DOI:** 10.1101/282699

**Authors:** Vadim Demichev, Christoph B. Messner, Spyros I. Vernardis, Kathryn S. Lilley, Markus Ralser

## Abstract

Data-independent acquisition (DIA-MS) boosts reproducibility, depth of coverage and quantification precision in label-free proteomic experiments. We present DIA-NN, a software that employs deep neural networks to distinguish real signals from noise in complex DIA datasets and a new quantification algorithm, that is able to subtract signal interferences. DIA-NN vastly outperforms the existing cutting-edge DIA-MS analysis workflows, particularly in combination with fast chromatographic methods, enabling deep and precise proteome coverage in high-throughput experiments.

## Introduction

Mass-spectrometry-based proteomics approaches involving artificial intelligence to dissect complex relationships between genotype and phenotype are rapidly gaining importance within both personalised medicine and the emerging field of data-driven biology^1–3^. These applications depend on large sample series and require reproducible and precise protein quantification. This is however hampered by the inherent complexity of the proteome, which leads to stochasticity in peptide detection via conventional data-dependent acquisition (DDA) strategies, resulting in missing values between successive runs^4,5^. Data-independent acquisition (DIA) approaches, such as SWATH-MS^6,7^, have been developed to reduce stochastic elements in proteomic data acquisition via sequential windowed fragmentation of all precursor ions (i.e. peptides with a specific charge) within a specified mass range. DIA workflows show high reproducibility and achieve superior proteomic depth, becoming the method of choice for protein identification and quantification in large sample series^8–10^. The computational processing of DIA datasets, however, is extremely challenging due to their inherent complexity. First, each precursor ion gives rise to a series of consecutive spectra in the data (instead of a single spectrum in a typical DDA workflow). Each of its fragment ions thus corresponds to an elution profile (i.e. a chromatogram). Second, mass-windowed acquisition leads to co-fragmentation of multiple interfering precursors (i.e. precursors that share some fragments with similar m/z values), leading to highly multiplexed spectra. Despite the numerous software improvements introduced recently^11^, only a fraction of the recorded information is currently efficiently extracted from the DIA data, hindering the identification performance. In addition, although quantification in DIA is performed at the MS^2^ level unlike most DDA methods, it is still affected by interferences that scale with increasing sample complexity or shorter chromatographic gradients, requiring more sophisticated algorithms to identify, or correct, for them^11^. In parallel, analysis of larger-scale DIA experiments is further limited by huge hardware demands or slow processing times of currently available software. Recently, new types of spectral deconvolution strategies aimed at better handling of signal interferences have been implemented in software tools such as Specter or microDIA^12,13^, but these still leave the requirements for the processing of large sample series, especially those acquired using fast chromatographic gradients, largely unaddressed.

## Results

We have developed and benchmarked DIA-NN, a software based on the novel application of deep neural networks to DIA data, which vastly outperforms the existing cutting-edge pipelines in both the identification numbers and quantification precision, as required for the next generation of high-throughput proteomics. The fully-automatic DIA-NN workflow (Figure 1A; all procedures are described in detail in Supplementary Methods), starts with a peptide-centric approach^14^, based on a spectral library, which can be provided separately or automatically generated by DIA-NN *in silico* from a protein sequence database (Supplementary Notes 2 and 3). First, a library of negative controls (i.e. decoy precursors^14,15^) is generated, to complement the library of real (i.e. target) precursors. For each target or decoy precursor, chromatograms are extracted from the raw DIA data and putative elution peaks (comprising the precursor and fragment ion elution profiles in the vicinity of the putative retention time of the precursor) are identified. A set of scores is then calculated to describe each of the elution peaks (in total, DIA-NN calculates 69 different peak scores in the various steps of the workflow). The scores reflect peak characteristics such as co-elution of fragment ions, mass accuracy or similarity between observed and reference (library) spectra (Supplementary Table 1 for details of the scoring system). The best candidate peak is then selected per precursor using iterative training of a linear classifier, which allows to calculate a single discriminant score for each peak.

While being highly sensitive^16^, the peptide-centric search alone leads to false identifications and unreliable quantification, as a single putative elution peak in the data can be used as the detection evidence for several precursors that share one or more fragments with close m/z values. DIA-NN tackles this by drawing upon the advantages of spectrum-centric approaches. It looks for all such situations when potentially interfering precursors have been matched to the same retention time (by the peptide-centric search module), and, if the degree of interference is deemed significant enough, only reports the ones best supported by the data as identified.

To calculate the precursor q-values, all target and decoy precursors need to be assigned a single discriminant score each, based on the characteristics of the respective candidate elution peaks. In DIA-NN, this crucial step in the workflow, which determines the number of precursors reported at a given false discovery rate (FDR) threshold, relies on deep neural networks (DNNs). DNNs encompass a group of artificial intelligence methods, that have been developed extensively in recent years, making them the preferred machine learning approach for many applications involving the analysis of complex data of heterogeneous nature^17^. Linear classifiers, conventionally used to score precursors, are unable to effectively deal with the highly complex DIA data. In DIA-NN, an ensemble of DNNs is trained to distinguish between the target and decoy precursors. For each precursor, the set of scores corresponding to the respective elution peak is used as neural network input. Subsequently, each trained network, when provided with a set of scores as input, yields a quantity that reflects the likelihood that this set originated from a target precursor. These quantities, calculated for all the precursors and averaged across the networks, are then used to obtain the q-values.

Furthermore, DIA-NN introduces an effective algorithm for detection and removal of interferences from tandem-MS spectra. For each putative elution peak, DIA-NN selects the fragment least affected by interferences (as the one with the elution profile best correlated with the elution profiles of the other fragments). Its elution profile is then considered representative of the true elution profile of the peptide. Comparison of this profile with the elution profiles of other fragments allows to subtract interferences from the latter.

In combination with the enhanced precursor scoring by DNNs, this new quantification strategy leads to a vast improvement of DIA data extraction, specifically in the analysis of complex proteomes with short chromatographic methods that suffer the most from the problem of signal interferences. To illustrate the performance of DIA-NN, we benchmarked it on the basis of public datasets that have been specifically created for testing DIA software. Its identification performance was evaluated using a HeLa whole-proteome tryptic digest recorded on a nanoLC-coupled QExactive HF mass spectrometer (Thermo Fisher), with different chromatographic gradient lengths, ranging from 0.5h to 4h^10^. The same data were processed with state-of-the-art DIA processing workflows currently used: OpenSWATH^18^, Skyline^19^ and Spectronaut^4^ (Biognosys). The number of precursor IDs produced by each tool is plotted as a function of the estimated effective FDR. This analysis demonstrated vastly better identification performance of DIA-NN, particularly evident at strict FDR thresholds and short chromatographic gradient lengths (Figure 1B; please see Supplementary materials for all peptide identification tables, which have been deposited online). At 1% estimated FDR, DIA-NN identifies more precursors from the 0.5h chromatographic gradient than Skyline or OpenSWATH from the same sample when analysed using a 2h chromatographic gradient. Thus, if a similar number of precursor identifications at 1% FDR is attempted, a change of software from either open-source tool to DIA-NN would allow sample throughput to be increased by a factor of ~four.

As misidentified peptides result in unreliable quantification, we next reasoned that the superior identification capabilities of DIA-NN would in itself lead to more precise quantification. To provide a minimally biased comparison, we only benchmarked DIA-NN against Spectronaut, which demonstrated the most similar identification performance on data collected using longer chromatographic gradients (Figure 1B). We used the LFQbench dataset created as part of a multi-center study and specifically designed to compare the quantification performance of DIA software tools^20^. In LFQbench, two peptide preparations (yeast and *E.coli*) were spiked in two different ratios (A and B) into a third a peptide preparation (human). The LFQbench reveals quantification precision on the basis of how well the ratios between the yeast, *E. coli* and human peptides (and proteins) are recovered. Of note, LFQbench data have been recorded on a TripleTOF 6600 mass spectrometer (Sciex), and serves hence also as a test of how well DIA-NN performs on different mass spectrometry platforms. DIA-NN demonstrated significantly better precision in the quantification of both yeast and *E.coli* peptides and proteins, while also generating more valid A:B peptide and protein ratios for each species (Figure 1C, Supplementary Figure 1). In addition, DIA-NN produced substantially better median CV values for human peptides and proteins: 5.4% and 2.9%, respectively, compared to 7.0% and 3.8% for Spectronaut, as calculated by the LFQbench R package.

In order to make DIA-NN accessible for the broad application in small-scale and large-scale proteomic experiments, we have included several additional features and programmed a comprehensive software tool that allows the conducting of all steps of a DIA-processing pipeline automatically. DIA-NN includes an intuitive graphical interface (screenshot in Supplementary Figure 4), as well as a command line tool for efficient integration into automated workflows. DIA-NN can analyse data generated on different mass spectrometry platforms, and does not require retention time standards to be present in the sample. DIA-NN also performs automatic mass correction and automatically determines such search parameters as the retention time window and the extraction mass accuracy. This eliminates the lengthy and laborious process of optimising the processing workflow for each particular data set. Moreover, written in C++, DIA-NN achieves ultrafast processing times with moderate hardware requirements, enabling fast and precise extraction of peptide and protein quantities from large-scale DIA proteomics datasets (100s – 1000s of samples) (Supplementary Note 4).

Finally, DIA-NN includes a software module that enables it to operate without a spectral library and generate high-quality spectral libraries directly from DIA data (Supplementary Note 2 an 3). The library-free mode is efficient for applications in which proteomic depth does not need to be exhausted and enables applications in which limited sample amount or instrument access restrictions prevent the creation of an extensive DDA-based spectral library^16^.

**Figure 1.**
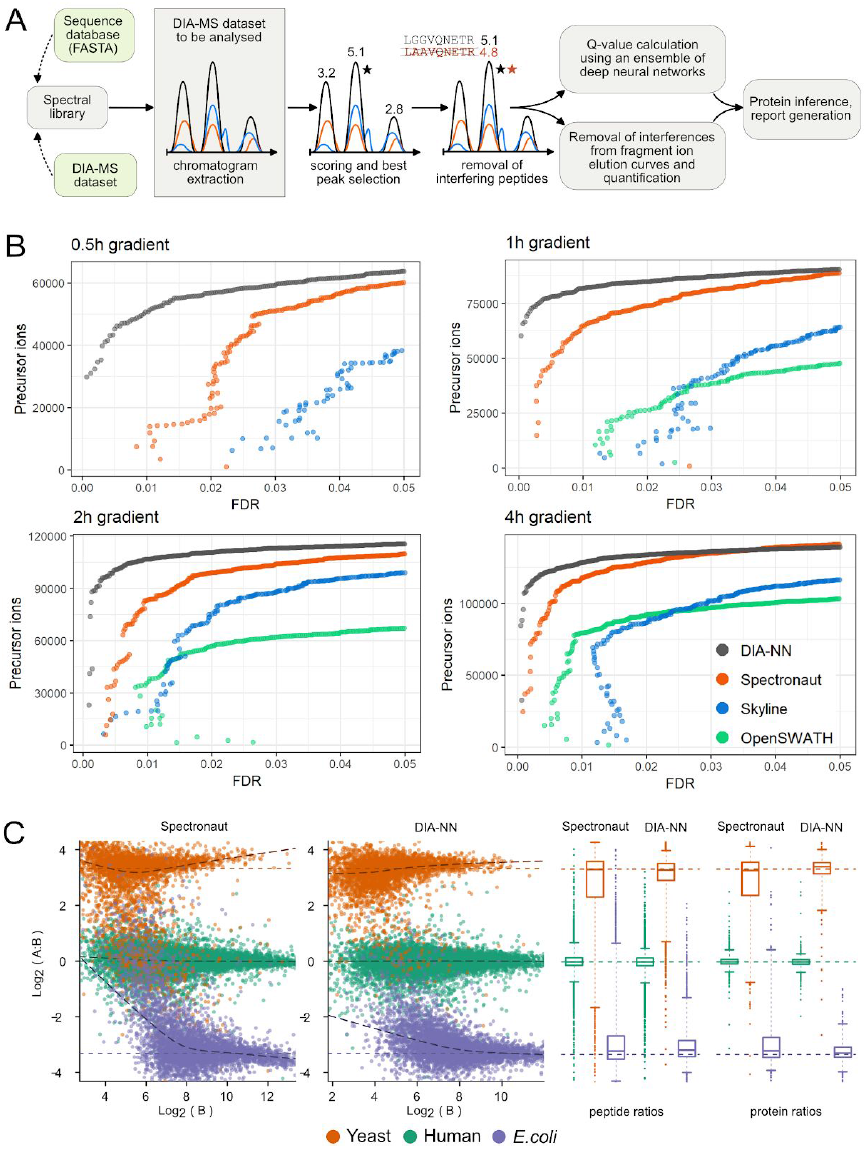
DIA-NN workflow and its performance. (A) DIA-NN workflow. (B) Identification performance of DIA-NN compared to OpenSWATH^18^ (part of OpenMS^21^ 2.3.0, released on January 3, 2018), Skyline^19^ (4.1.0.11796, released on January 11, 2018) and Spectronaut^4^ (Pulsar 11.0.15038.17.27438 (Asimov) (Biognosys), released on June 2, 2017).

The benchmark illustrates the results of processing the raw data files generated from the analysis of a HeLa cell-line whole-proteome tryptic digest recorded on a nanoLC-coupled QExactive HF mass spectrometer with chromatographic gradient lengths ranging from 0.5h to 4h^10^. OpenSWATH was not used to analyse the 0.5h run, as it was unable to correctly recognise the iRT retention time standards in the short gradient (Biognosys). The effective false discovery rate (FDR) was estimated using a two-species compound spectral library method^10^, i.e. a concatenated spectral library containing precursor ions mapped to either human or maize proteomes was used, the maize precursors serving as externally supplied decoys (as in a target-decoy method^15^) (see Methods). Each point on the graph corresponds to a decoy (maize) precursor, its x-axis value reflecting the estimated FDR at the respective score threshold and its y-axis value being the number of identified target (human) precursors at this threshold. DIA-NN consistently outperforms the other software tools in terms of the identification performance, in particular on short chromatographic gradients and with conservative FDR-thresholds. (C) Quantification precision, benchmarked using the LFQbench test performance of DIA-NN in comparison to Spectronaut, in the analysis of peptide preparations (yeast and *E.coli*) that were spiked in two different proportions (A and B) into a human peptide preparation^20^. The data were processed at 1% q-value, and peptide ratios between the mixtures were visualised using the LFQbench R package (with the dotted lines indicating the expected ratios). Right panel: peptide and protein quantification performance given as box-plots. DIA-NN demonstrates significantly better quantification precision for both yeast and *E.coli* peptides.

## Acknowledgements

We thank Roland Bruderer (Biognosys) for providing the spectral libraries. This work was supported by the Francis Crick Institute which receives its core funding from Cancer Research UK (FC001134), the UK Medical Research Council (FC001134), and the Wellcome Trust (FC001134), and received specific funding from the BBSRC (BB/N015215/1 and BB/N015282/1), as well as a Crick Idea to Innovation (i2i) initiative (Grant Ref 10658).

## Methods

### Raw mass spectrometry data

Raw analyses of the HeLa cell lysate have been described previously^1^ and were obtained from ProteomeXchange (data set PXD005573). DIA-NN and Spectronaut accessed these directly; for processing with Skyline and OpenSWATH, .raw files were converted to the .mzML format using MSConvertGUI (part of ProteoWizard^2^ 3.0.11537) with MS1 and MS2 vendor peak picking enabled, 32-bit binary precision and all other options unchecked. Raw data files for the LFQbench test were generated by Navarro and colleagues^3^ and were obtained from ProteomeXchange (data set PXD002952; HYE110 runs on TripleTOF 6600 with 64-variable windows acquisition). For the analysis with DIA-NN, these were converted to the .mzML format using MSConvertGUI.

### Spectral library

The human and maize spectral libraries used to generate the two-species compound library have been described previously^1^. The maize library was filtered to exclude peptides matched to either the NCBI human redundant database (April 25^th^, 2018) or the UniProt^4^ human canonical proteome (3AUP000005640). The human library was filtered to include only peptides matched to the latter. In both cases, filtering was performed with leucine and isoleucine treated as the same amino acid. The libraries were merged, resulting in a library containing only precursor ions matched to either human or maize proteomes, but not both. To enable the use of the library by all of the software tools under consideration, the library was converted to the OpenMS-compatible format with the use of DIA-NN. Following the protocol of Navarro and co-workers^3^, only precursor ions associated with at least six fragment ions were retained in the library, and all fragments but the top six (ordered by their reference intensities) were discarded. This was done to ensure that there is no bias in terms of the distribution of the number of annotated fragments between human and maize precursors. In addition, although DIA-NN can take advantage of large numbers of fragment ions described in the spectral library, many software tools tend to perform poorly if the number of fragment ions is not restricted, e.g. Spectronaut and Skyline only use the top six fragments by default. Reference retention times (Biognosys iRT scale) below ‐60.0 were adjusted to ‐60.0, to enable efficient linear retention time prediction by Skyline and OpenSWATH, as the respective precursors were observed to elute concomitantly. A low number of precursor ions had to be removed from the spectral library, so that the library could be imported error-free into Skyline (Supplementary Table 2). The resulting compound spectral library contained 202310 human precursor ions and 9781 maize precursor ions.

### FDR estimation using the compound spectral library

The HeLa cell lysate proteomic datasets (Figure 1B) were analysed with each software tool using the human-maize compound spectral library described above. For each identified maize precursor, its score (that was ultimately used to calculate the q-value) was considered. The numbers of human and maize precursors identified with the same or better score were then calculated ([human IDs] and [maize IDs], respectively). A conservative FDR estimate was then obtained:

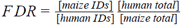

Here [human total] and [maize total] are the respective numbers of human and maize precursors in the spectral library.

### Configuring DIA-NN and Spectronaut

The default settings were used for DIA-NN (v1.5.7) and Spectronaut, except that protein inference and FDR filtering of the output were turned off to obtain complete reports.

### Configuring Skyline and OpenSWATH

For Skyline and OpenSWATH, we used the default settings and settings described previously^3^ whenever possible. The spectral library was directly imported into Skyline using the 0.05 m/z ion match tolerance. Shuffle decoy generation was used. For the use with OpenSWATH, the spectral library was converted (using OpenMS 2.3.0) to the .TraML format, decoys were generated using the following options: “-append ‐exclude_similar ‐remove_unannotated ‐enable_detection_specific_losses ‐enable_detection_unspecific_losses-force”. The spectral library was then converted back to the .tsv format. Several mass accuracy settings were attempted (separately for each run); in each case the setting yielding the highest number of reported precursor ion identifications at 1% q-value was chosen: 7ppm for 0.5h, 1h and 2h, 5ppm for 4h – Skyline, 15ppm for 1h and 2h, 10ppm for 4h – OpenSWATH. For all runs but the 4h run, the retention time window was set to 20 minutes (Skyline) and 10 minutes (OpenSWATH). For the 4h run, the retention time window was set to 40 minutes (Skyline) and 20 minutes (OpenSWATH). Skyline was run with the acquisition method set to DIA, product mass analyzer set to centroided and isolation scheme set to “Results (0.5 margin)”. (We also attempted running Skyline with product mass analyzer set to Orbitrap on the 2h and 4h gradient .raw files without converting to centroided .mzML, but this resulted in a significantly lower number of identified precursors.) The retention time calculator was created using the “Biognosys-11” built-in set of retention time standards. The calculation of q-values was performed using the built-in mProphet algorithm. OpenSWATH was run using the following options in addition to setting the mass accuracy and the retention time window:

“-readOptions cacheWorkingInMemory ‐batchSize 1000

-Scoring:TransitionGroupPicker:background_subtraction original

-Scoring:stop_report_after_feature ‐1 ‐Scoring:Scores:use_dia_scores true ‐ppm ‐threads 12

-min_upper_edge_dist 1.0 ‐min_rsq 0.95 ‐tr_irt iRTassays.TraML

-extra_rt_extraction_window 100 ‐use_ms1_traces”.

The “-min_coverage” option was set to 0.5 for 1h and 0.6 for 2h and 4h runs. Retention time standards for OpenSWATH were provided in the iRTassays.TraML file downloaded from the PeptideAtlas^5^ repository with the identifier PASS00779. OpenSWATH output was further processed using PyProphet^6^ 2.0.0 with the “‐‐level=ms2” option. PyProphet output was further processed in R to remove decoy precursors and suboptimal peaks.

## Supplementary Notes

### 1. Performance of DIA-NN in the LFQbench test

While the identification performance is important, so far the key application of DIA is accurate, precise and consistent peptide and protein quantification in large sample series. We illustrated the quantification performance of DIA-NN by comparing it to Spectronaut Pulsar using the LFQbench test^1^ (HYE110 dataset, 64 variable window acquisition scheme on TripleTOF 6600) (Supplementary Figure 1). In this benchmark, human, yeast, and *E.coli* lysates were mixed in different proportions and analysed via SWATH-MS. For each mixture, three injection replicates were measured. The performance of the software tools was compared using the LFQbench R package (https://github.com/IFIproteomics/LFQbench), which takes as input the intensities of the precursor ions and uses these to quantify peptides and proteins. Q-value threshold was set to 1%. The default settings were used for DIA-NN and Spectronaut, except that protein inference and FDR filtering of the output were turned off to obtain complete reports.

**Supplementary Figure 1.**
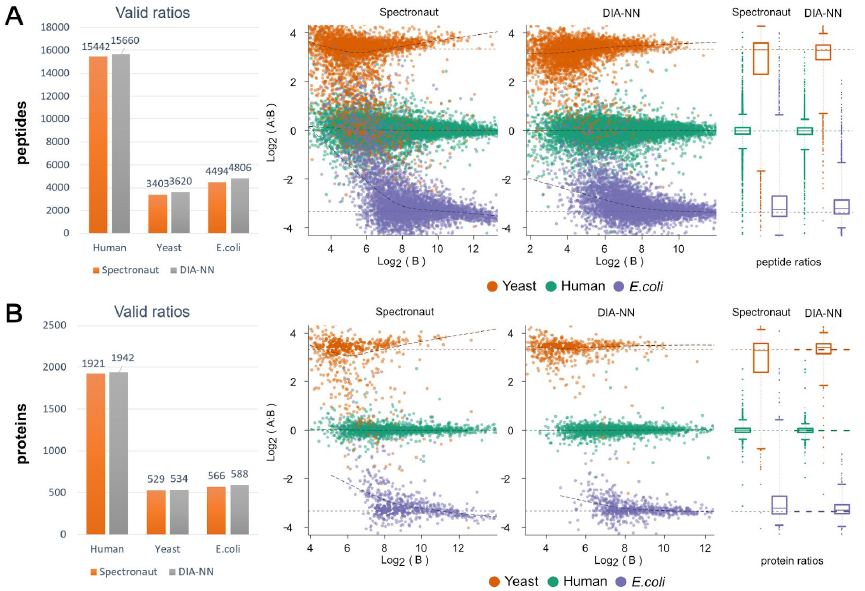
Performance of DIA-NN in the LFQbench test (Complete Figure, of which an extract is shown in Figure 1C). LFQbench performance of DIA-NN in comparison to Spectronaut. In the LFQbench test, two peptide preparations (yeast and *E.coli*) are mixed in two different proportions (A and B), pooled with a human peptide preparation and analysed in triplicates on TripleTOF 6600^1^. The data were processed at 1% precursor q-value; peptide (panel A) and protein (panel B) ratios between the mixtures were visualised using the LFQbench R package (with the dotted lines indicating the expected ratios). DIA-NN demonstrates significantly better quantification precision for both yeast and *E.coli* peptides and proteins, as evidenced by the box plots for the ratios. DIA-NN also produced better median CV values for human peptides and proteins: 5.4% and 2.9%, respectively, compared to 7.0% and 3.8% for Spectronaut, as calculated by the LFQbench R package.

### 2. Library-free processing

DIA-NN can process raw data using either a spectral library or a protein sequence database. In the latter case, proteins are *in silico* digested and prediction of the fragmentation spectra of the resulting peptides as well as the respective retention times is performed (Supplementary Methods). While the library-based approach achieves higher proteomic depth, the library-free approach saves sample material, as well as the instrument time. We benchmarked the library-free performance of DIA-NN using the HeLa proteome raw data with different chromatographic gradient lengths^2^ (Supplementary Figure 2). Library-free analysis was carried out against the human UniProt canonical proteome (3AUP000005640) with the maximum peptide length set to 50 and a single missed cleavage allowed. To demonstrate the benefit of restricting the search space, the data were also processed with the same sequence database but filtered to include only peptides known to be present in human samples, according to the PeptideAtlas^3^ build of January 2018; for this, the maximum peptide length was set to 100 and up to five missed cleavages were allowed. An *E.coli* spectral library^2^ was used to train the peptide fragmentation and retention time predictors. A project-specific spectral library^2^ was used for library-based processing.

**Supplementary Figure 2.**
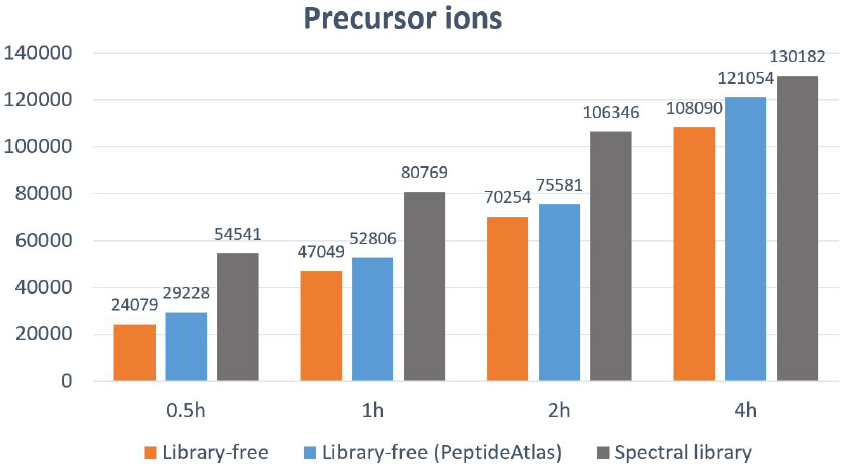
Library-free performance of DIA-NN. The numbers of precursors identified at 1% q-value threshold, as reported by DIA-NN, as a function of chromatographic gradient length^2^ (h).

### 3. Generating spectral libraries with DIA-NN

DIA-NN can generate spectral libraries directly from DIA data. Here, we demonstrate its capabilities using a workflow optimised for high-throughput proteome quantification based on 23-minute gradient microflow SWATH^4^ applied on yeast and human plasma proteomes (Supplementary Figure 3). Briefly, Sciex TripleTOF 6600 was used to rapidly analyse yeast and human plasma tryptic digests (three injections each; see the detailed workflow description in the Supplementary Methods section). Furthermore, a set of SWATH gas-phase fractionation runs with narrow precursor isolation windows was acquired for each of the digests. DIA-NN was used to create DIA-based spectral libraries directly from these gas-phase fractionation runs. For the yeast library, a search against the yeast UniProt canonical proteome was used (3AUP000002311). For the plasma library, the runs were searched against the human UniProt canonical proteome (3AUP000005640) filtered for the peptides known to be present in human plasma, according to the PeptideAtlas^3^ build of August 2013; for this, the maximum peptide length was set to 100 and up to five missed cleavages were allowed. The numbers of proteins uniquely identified at 1% q-value (i.e. using peptides specific to the respective genes) were then calculated, as well as the numbers of these with coefficients of variation (CV) less than the specified thresholds (measured for ubiquitously identified proteins). Yeast runs were also analysed by DIA-NN directly, without the DIA-based spectral library. For the initial analysis of yeast gas-phase fractionation runs, an *E.coli* spectral library^2^ was used to train the peptide fragmentation and retention time predictors. Subsequently, all library-free processing was performed using this library, specific to our LC-MS setup, to train the predictors. The 1% precursor q-value threshold was used for all the analyses.

**Supplementary Figure 3.**
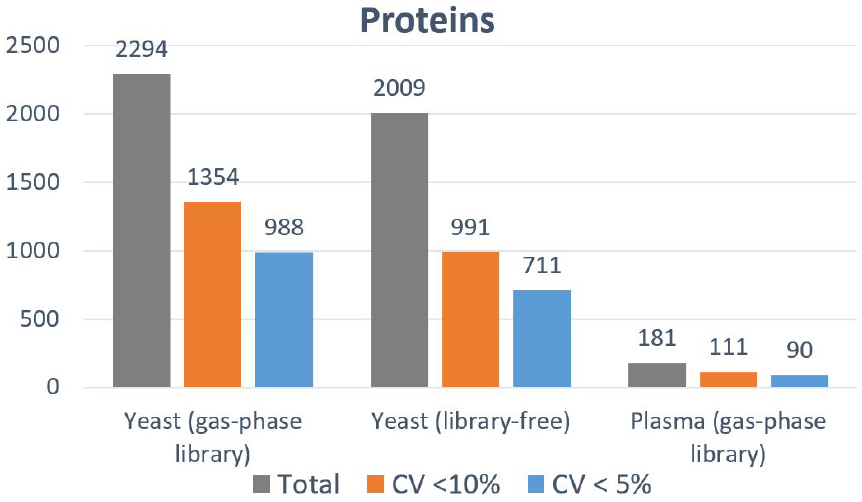
Using DIA-NN to analyse yeast and human plasma SWATH runs without a DDA-based spectral library. DIA-NN was used to analyse yeast and human plasma triplicate 23-minute gradient runs using spectral libraries generated by DIA-NN from gas-phase fractionation runs. A library-free analysis of the same yeast runs was added for comparison. Only uniquely identified proteins (i.e. using proteotypic peptides only) were considered (filtered at 1% precursor-level and 1% protein-level q-value).

### 4. Hardware requirements, speed and GUI

The rising interest in high-throughput proteomics in research, medicine and industry calls for the development of software tools that are able to rapidly and reliably analyse thousands of mass spectrometry runs. DIA-NN performs the computationally-demanding processing steps separately for each run in the experiment, saving all the relevant information to compact files on the hard drive. This allows quick and flexible analysis and subsequent reanalysis of any part of the experiment separately. In addition, DIA-NN is very fast. For example, on an average workstation (2x 6-core Xeon E5645 @2.4Ghz), it required less than 12 minutes to analyse the four HeLa runs (used to generate Figure 1B) and less than 17 minutes to analyse the three yeast runs (used to generate Supplementary Figure 3) in library-free mode. Finally, it has very low hardware requirements, e.g. during the processing of the HeLa and yeast proteomes its memory usage peaked at less than 3.3Gb and 1.8Gb, respectively.

For the large scale applications, we provide a command line tool for the creation of automatic processing workflows. For smaller or more routine applications, we have further programmed a graphical user interface (GUI) wrapper, that enables the control of all steps of the workflow from a simple and intuitive workspace (Supplementary Figure 4). Although DIA-NN is designed to do as much as possible automatically, it is fully configurable, allowing to fine-tune the processing workflow for a specific experiment. The GUI allows to easily set up the analysis in few clicks without losing the powerful tuning capabilities of the command line tool.

**Supplementary Figure 4.**
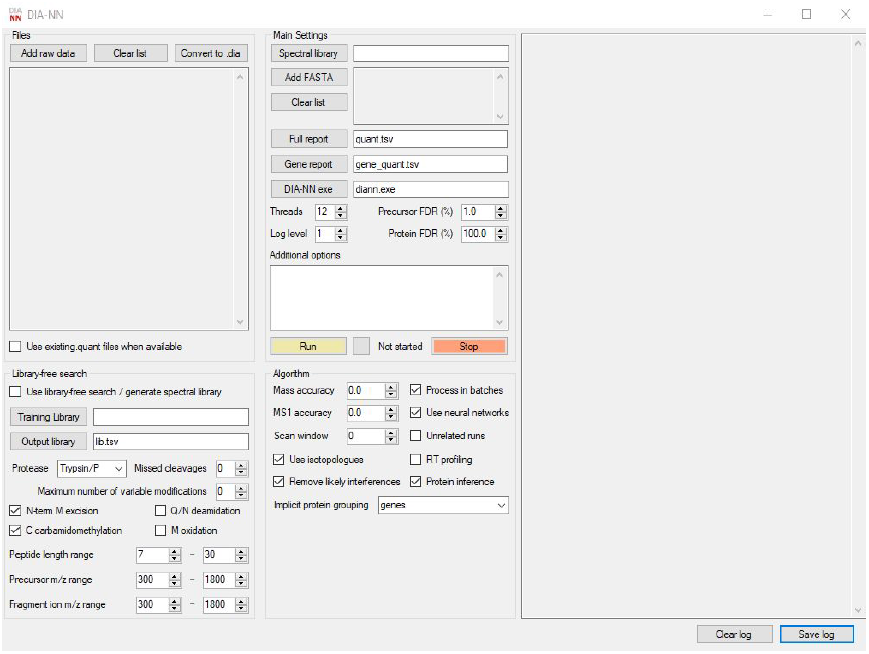
DIA-NN graphical user interface.

### 5. Supplementary Methods

#### DIA-NN algorithms

DIA-NN is a fully open-source software tool and is freely available at https://github.com/vdemichev/diann. Here we describe the algorithms used by DIA-NN.

DIA-NN requires either a spectral library or a sequence database to be provided as input. In the latter case, DIA-NN generates a spectral library *in silico*. For this, DIA-NN can optionally use a fragmentation predictor (based on the approach introduced in MS Simulator^5^) and a linear retention time predictor. The predictors are trained using any spectral library supplied by the user.

For each target precursor in the spectral library, a decoy precursor is generated, if not provided in the library. By default this is done by replacing the fragment ion m/z values of the target precursor assuming the amino acids adjacent to the peptide termini were mutated (GAVLIFMPWSCTYHKRQEND to LLLVVLLLLTSSSSLLNDQE mutation pattern is used). Optional pseudo-reverse approach to decoy precursor generation is also supported.

Chromatograms are then extracted for each target and decoy precursor and the respective fragment ions. Potential elution peaks are identified, and for each of these the fragment with the most optimal properties for quantification is selected. This fragment (chosen among the top six based on the reference intensities in the library) maximises the sum of the Pearson correlations between its elution profile (in the vicinity of the putative peak) and the elution profiles of the remaining fragments from the top six list. It is assumed, that this “best” fragment is likely to be the one least affected by interferences, its elution profile thus being representative of the true elution profile of the peptide. A set of 69 scores is calculated for each potential elution peak (Supplementary Table 1). These are used differentially in different processing stages based on algorithmic decision making. The “best” candidate peak is selected per precursor using one of the scores, and a linear classifier is trained to distinguish between target and decoy precursors based on the sets of scores corresponding to the respective best peaks, allowing to calculate a single discriminant score for each peak. The discriminant scores are used to refine the selection of best peaks, and the procedure is repeated iteratively several times.

During the next step, DIA-NN looks for precursors matched to the same retention time which also have interfering fragments. If the degree of interference is deemed significant enough, DIA-NN only reports the precursor with the highest discriminant score as identified. This method effectively allows to combine the advantages of peptide-centric and spectrum-centric approaches to mass-spectrometry data analysis.

An ensemble of deep feed-forward fully connected neural networks (12 by default) is trained (as implemented in the Cranium library (https://github.com/100/Cranium) supplied with the DIA-NN distribution) via Adam^6^ to distinguish between target and decoy precursors. For each precursor, the set of scores corresponding to the respective best elution peak is provided as input for the networks. Each network comprises a series of *tanh* hidden layers (5 by default) and a softmax output layer. Cross-entropy is used as the loss function. The peak scores (69 total) are standardised before training. By default, training is performed for one epoch only, minimising the effects of overfitting. The predictions of the neural networks are then averaged for each precursor, resulting in the final set of scores used for q-value calculation. Optionally, DIA-NN can train each network on a part of the dataset, only using it to score precursors it has not been trained on, or use a higher number of training epochs. The use of neural networks allows to effectively utilise all the 69 scores calculated for each elution peak, thus increasing the amount of information extracted from the data in comparison to the use of a linear classifier.

For a particular score threshold, DIA-NN calculates a conservative FDR estimate (used to generate the respective q-values^7^ for precursor identifications), dividing the number of decoys with scores exceeding the threshold by the number of targets with scores exceeding the threshold. Correction based on estimating the prior probability of incorrect identification (π_0_) is not performed.

DIA-NN uses a conservative protein q-value calculation method, which is applied to individual proteins and not protein groups. To estimate protein-level FDR, only target and decoy precursors specific to the protein of interest are considered. Thus, proteins without any proteotypic precursors identified are automatically assigned a q-value equal to one. The maxima of target and decoy scores are calculated for each protein and the distributions of these are examined. For a given score threshold, FDR is estimated by dividing the number of decoy scores exceeding it by the number of target scores exceeding it.

For each run, DIA-NN quantifies the intensities of all fragment ions associated with each precursor. For this we have conceived an efficient interference removal algorithm. The elution profile *x* (·) of each fragment is compared to the reference profile *ref* (·), the smoothed elution profile of the best fragment (the one defined previously for the potential elution peak being considered). The “weighted” fragment intensity is calculated as the sum of the fragment elution profile values weighted by the respective squared values of the reference profile. This emphasises the contribution of the data points close to the apex of the reference elution profile, thus making the impact of potential interferences manifesting far from the apex negligible. The ratio *r* of weighted intensities of the fragment under consideration and the best fragment is calculated. All values of *x* (·) exceeding 1.5·*r* ·*ref* (·) are replaced with 1.5·*r* ·*ref* (·). The area under the resulting profile is then considered to be the intensity of the fragment. Preliminary precursor quantities are obtained by summing the quantities of the top six fragments (ranked by their library intensities).

DIA-NN enables cross-run precursor ion quantification. In each run, each fragment is assigned a score which is the correlation score of its elution profile with the respective reference profile, i.e. the smoothed elution profile of the best fragment. For each precursor, three fragments with highest average correlations are selected in a cross-run manner. Only runs where the precursor was identified with a q-value below a given threshold are considered. The intensities of these fragments are then summed in each run to obtain the precursor ion intensity. This approach allows to deal with the situation when in certain runs interferences were not efficiently removed from elution profiles of some fragments, e.g. if interferences manifested close to the apexes.

Protein grouping can be performed either for individual runs or in a cross-run fashion (default). For each precursor, DIA-NN aims to reduce the number of proteins associated with it using the maximum parsimony principle, which is implemented via a greedy set cover algorithm.

After precursor ion quantification, cross-run normalisation and protein quantification are performed. All the precursor intensities corresponding to identifications with q-values above a given threshold are replaced with zeros and preliminary cross-run normalisation based on the total signal (i.e. the sum of the intensities of all precursors) is performed. Precursors are then ordered by their coefficients of variation. Top *pN* precursors are selected, where *N* is the average number of identifications passing the q-value threshold and *p* is between 0 and 1. Sums of the intensities of these precursors are calculated and are used for normalisation, i.e. the levels of all precursors are scaled to make these quantities equal in different runs. A “Top 3” method is eventually used for protein quantification: intensities of protein groups are calculated as sums of the intensities of top 3 most abundant precursors identified with a q-value lower than a given threshold in a particular run.

#### Sample preparation and mass spectrometry

The yeast protein extracts were prepared from *Saccharomyces cerevisiae* (BY4743-pHLU^8^) grown to exponential phase in minimal synthetic nutrient media and processed in a bead beater for 5min at 1500rpm (Spex Geno/Grinder). Plasma samples were prepared from commercial plasma (Human Cord Blood Plasma, Stemcell Technologies).

Proteins were denatured in 8M urea/0.1M ammonium bicarbonate pH 8.0 before they were reduced and alkylated in 5mM dithiothreitol and 10mM iodoacetamide, respectively. The sample was diluted to <1.5M urea/0.1M ammonium bicarbonate pH 8.0 before the proteins were digested overnight with trypsin (37°C). Peptides were cleaned-up with 96-well MacroSpin plates (Nest Group) and iRT peptides (Biognosys) were spiked in.

The digested peptides were analysed on a nanoAcquity (Waters) coupled to a TripleTOF 6600 (Sciex). Peptides were separated with a 23-minute non-linear gradient (4% acetonitrile/0.1 % formic acid to 36% acetonitrile/0.1% formic acid) on a Waters HSS T3 column (150mm x 300µm, 1.8µm particles) with a 5µl/min flow rate. The DIA method consisted of an MS1 scan from m/z 400 to m/z 1250 (50ms accumulation time) and 40 MS2 scans (35ms accumulation time) with variable precursor isolation width covering the mass range from m/z 400 to m/z 1250.

The library generation with “gas-phase fractionation” was performed using the same LC-MS/MS setup as mentioned above. The peptides were separated with a 120 minute (plasma samples) and 45 minute (yeast samples) linear gradient (3% acetonitrile/0.1% formic acid to 60% acetonitrile/0.1 formic acid). Repetitive injections were performed to cover the following scan ranges: m/z 400 – 500, m/z 495 – 600, m/z 595 – 700, m/z 695 – 800, m/z 795 – 900, m/z 895 – 1000, m/z 995 – 1100, m/z 1095 – 1250 (yeast) and m/z 400 – 500, m/z 500 – 600, m/z 600 – 700, m/z 700 – 800, m/z 800– 900, m/z 900 – 1000, m/z 1000 – 1250 (plasma). The precursor selection windows were m/z 4 (m/z 1 overlap) for all runs except the yeast m/z 1095 – 1250, for which m/z 5 (m/z 1 overlap) windows were used. For the plasma runs, each acquisition cycle was split into two subcycles with the second subcycle having the isolation windows shifted by m/z 1.5.

**Supplementary Table 1.** Scoring of putative elution peaks by DIA-NN. Groups of scores (69 in total) calculated by DIA-NN for all putative elution peaks (matched to the respective target or decoy precursor ions).

| **Score group** | **Number of scores** |
| --- | --- |
| Ions co-elution (MS2 level) | 18 |
| Ions co-elution (MS1 level) | 3 |
| Isotopologue ions co-elution | 11 |
| Total signal | 1 |
| Measured relative fragment intensities | 7 |
| Mass accuracy (MS2) | 6 |
| Retention time (RT) | 2 |
| Elution profile shape | 5 |
| Presence of other putative elution peaks | 2 |
| Library characteristics of the precursor | 14 |

**Supplementary Table 2.**
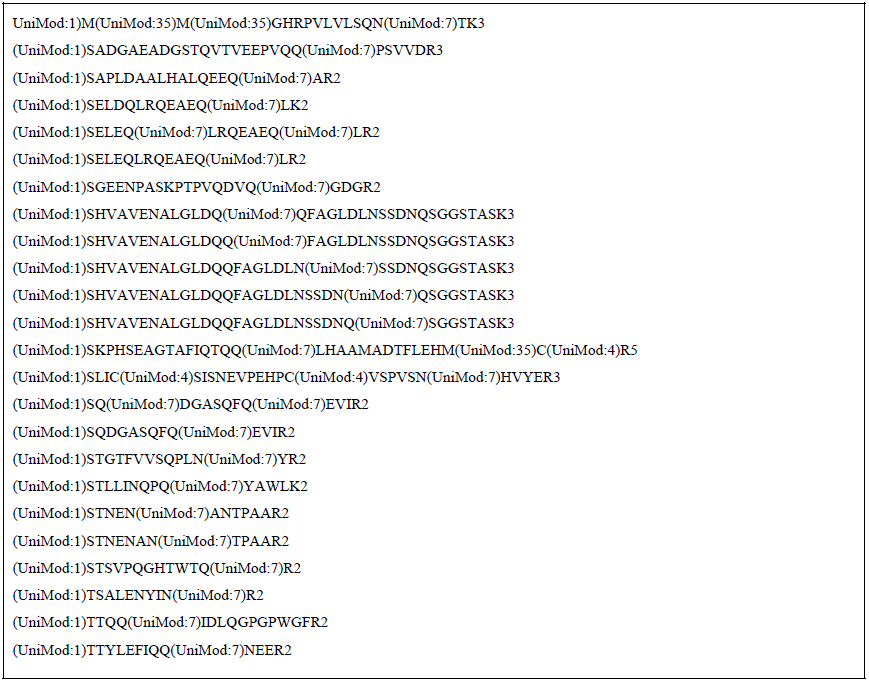
Precursor ions removed from the human-maize concatenated spectral library. Peptide modifications are encoded in the UniMod format, precursor charges are indicated with a number following the amino acid sequence.

